# Neurocognitive development of inhibitory control and substance use vulnerability

**DOI:** 10.1101/781286

**Authors:** Alina Quach, Brenden Tervo-Clemmens, William Foran, Finnegan J. Calabro, Tammy Chung, Duncan B. Clark, Beatriz Luna

**Author notes:** Equal Contribution (Alina Quach, Brenden Tervo-Clemmens). Corresponding Author: Brenden Tervo-Clemmens, M.S.

## Abstract

Previous research indicates that risk for substance use is associated with poor inhibitory control. However, it remains unclear whether at risk youth use follow divergent patterns of inhibitory control development. As part of the longitudinal National Consortium on Adolescent Neurodevelopment and Alcohol (NCANDA) study, participants (*N* = 113, baseline age: 12-21) completed a rewarded antisaccade task during fMRI, with up to three time points. We examined whether substance use risk factors, including dimensional measures of psychopathology (externalizing, internalizing) and family history of substance use disorder, were associated with developmental differences in inhibitory control performance and BOLD activation at both the trial-level and within individual antisaccade epochs (cue, preparation, and response). Among substance use risk factors, only externalizing psychopathology predicted developmental differences in inhibitory control, where high externalizing predicted lower correct response rates and faster latencies were observed in early adolescence, but normalized by late adolescence. Neuroimaging results revealed high externalizing was associated with developmentally-stable hypo-activation in the left middle frontal gyrus (trial-level), but divergent developmental patterns of posterior parietal cortex activation (cue epoch). Developmental differences in inhibitory control associated with externalizing suggest early adolescence may be a unique period of substance use vulnerability via cognitive and phenotypic disinhibition.

**Highlights:**

1. Characterized inhibitory control development in adolescents at-risk for substance use.
2. Externalizing psychopathology is associated with lower antisaccade correct response rate and faster latencies in early adolescence.
3. Externalizing performance differences normalize by late adolescence.
4. Externalizing is associated with prefrontal hypo-activation across development.
5. Externalizing moderates age-related increases in posterior parietal cortex activity.

## 1. Introduction

Across the lifespan, problematic substance use has been linked to impairments in inhibitory control: the ability to regulate prepotent behaviors, in order to facilitate goal-directed behaviors (Goldstein & Volkow et al., 2011). However, substance use initiation and escalation typically begin during adolescence (Johnston et al., 2018), when inhibitory control is still developing and ongoing brain maturation and specialization support the transition to adulthood (Larsen & Luna 2018; Ordaz et al., 2013). To this end, it has been suggested that diverging developmental trajectories of inhibitory control and other higher-order cognitive functions may increase vulnerability to problematic substance use (Casey et al., 2008, Steinberg, 2005), by way of early substance use experimentation (Johnston et al., 2018) and individual differences in psychiatric symptoms (Paus et al., 2008) that frequently co-occur with problematic substance use (Kreuger et al., 2002; Hussong et al., 2011). Supporting this perspective, recent work from our group has demonstrated that substance use risk factors, specifically externalizing psychopathology and impulsivity, are associated with poor inhibitory control and cortical activation differences among adolescents during an antisaccade task (Tervo-Clemmens, et al., 2017). However, it remains unclear whether certain periods of development are more sensitive to inhibitory control limitations associated with substance use vulnerability and if at-risk youth follow different trajectories of neurocognitive development.

Neuroimaging studies suggest that the protracted development of inhibitory control during adolescence relies on functional changes in brain regions supporting cognitive control across the lifespan, including dorsolateral prefrontal cortex, posterior parietal cortex, and the anterior cingulate cortex (Luna et al., 2015). Supporting a potential role of these regions in vulnerability to problematic substance use, poor response inhibition and reduced activation within the prefrontal cortex during inhibitory control tasks have been shown to predict problematic substance use in adolescence (Norman et al., 2011; Nigg et al., 2006). Similarly, prior work demonstrates adolescents with a family history of substance use disorders, who have an increased risk for problematic substance use (Cloninger, Sigvardsson, Reich, & Bohman, 1986) exhibit divergent age-related activation differences, relative healthy controls, within brain regions supporting inhibitory control, including the middle cingulate and frontal gyrus (Hardee et al., 2014). However, the field has yet to evaluate such relationships within the context of broader substance use risk factors, including dimensional measures of trait-level psychopathology, which prevents a comprehensive understanding of the developmental etiology of problematic substance use.

Leveraging a large longitudinal sample that performed an antisaccade inhibitory control task during fMRI acquisition as part of the National Consortium on Alcohol and Neurodevelopment in Adolescence (NCANDA) study, the current project examined whether substance use risk factors, including externalizing and internalizing psychopathology and family history of substance use disorder, were associated with deviations from normative neurocognitive development. Based on previous work implicating poor response inhibition in externalizing psychopathology, a latent construct characterized by behavioral undercontrol and impulsivity (Iacono, et al., 2008), and prior work examining non-developmental main effects of externalizing and impulsivity within a subset of the current sample (Tervo-Clemments et al., 2017), we predicted that externalizing would be associated with poor inhibitory control and reduced activation in regions supporting cognitive control, including the prefrontal cortex. Critically however, and novel to the current project, we further hypothesized that at-risk youth would follow developmental trajectories of inhibitory control that diverged from normative patterns, as would be predicted by current theories suggesting that neurodevelopmental limitations in cognitive control may underlie adolescent substance use vulnerability (Casey & Jones, 2010).

## 2. Methods

### 2.1 Participants

Participants were part of the University of Pittsburgh Medical Center Site of the NCANDA study. See Brown et al. (2015) for complete study protocol. Briefly, an accelerated longitudinal design was used to study adolescent brain development starting at various baseline ages (12 −21 years of age) with longitudinal follow ups. At the time of manuscript preparation, data collection included up to three time points (i.e., baseline and two annual follow-ups). In the interest of studying substance use risk and eventual initiation and escalation, the NCANDA study targeted high-risk youth with limited substance use. The study aimed to achieve a sample that consisted of approximately 50% of participants at risk for substance use, based on risk factors discussed below. Exclusion criteria included contraindications to magnetic resonance imaging (MRI; e.g., non-removable metal implements), medical history that would compromise MRI (e.g., traumatic brain injury), and a current or persistent major psychiatric disorder that would interfere with testing (e.g., psychosis). The study protocol was approved by the University of Pittsburgh Institutional Review Board. Informed consent and assent were obtained from adults and minors, respectively. Participants were compensated for their participation.

After excluding testing sessions (subject at visit) based on study-specific criteria (see below), the final sample for the current project included 113 participants (baseline age ranged from 12.27 - 21.96, M = 17.11, SD = 2.66). For the behavioral analysis, a total of 220 sessions were included. Among the sample, 41.6% (n = 47) of subjects had data which met criteria for baseline and follow-up and 26.5% (n =30) had data which met criteria at all three time points. The final neuroimaging sample included 104 participants and a total of 183 sessions, with 45.6% (n = 47) of the sample having data at both baseline and follow-up and 15.4% (n = 16) of the sample having data across all three visits (see below for neuroimaging exclusion criteria). See Figure 1 for sample age distribution and Table 1 for sample characteristics.

**Table 1.** Sample characteristics

|  |  | Baseline | Second Visit | Third Visit |
| --- | --- | --- | --- | --- |
| *N* |  | 94 | 79 | 47 |
| Female (%) |  | 59.57 | 53.16 | 44.68 |
| Age<sup>a</sup> |  | 17.11 (2.66) | 18.46 (2.48) | 19.09 (2.51) |
| Generalized Cognitive Ability (z-score)<sup>a, b</sup> |  | -0.01 (0.85) | 0.03 (0.87) | 0.25 (0.76) |
| Socioeconomic Status<sup>a</sup> |  | 89.63 (14.11) | 88.41 (15.32) | 91.36 (13.68) |
| Race (%) | Caucasian | 82.98 | 82.28 | 91.49 |
|  | Non-Caucasian | 17.02 | 17.71 | 8.51 |
| Substance use | Exceeds Threshold Drinking<sup>b</sup> (*n*) | 24 | 20 | 8 |
| Risk Factors | Externalizing T-score<sup>a, c</sup> | 42.91 (8.14) | 43.97 (8.71) | 41.49 (8.81) |
|  | Internalizing T-score<sup>a, c</sup> | 45.02 (9.41) | 45.25 (9.55) | 41.57 (10.36) |
|  | Family history of substance use disorder<sup>b</sup> (*n*) | 16 | 18 | 9 |

**Note.** <sup>a</sup>Mean (standard deviation). <sup>b</sup>Defined at baseline. <sup>c</sup>Gender and age adjusted

**Figure 1.**
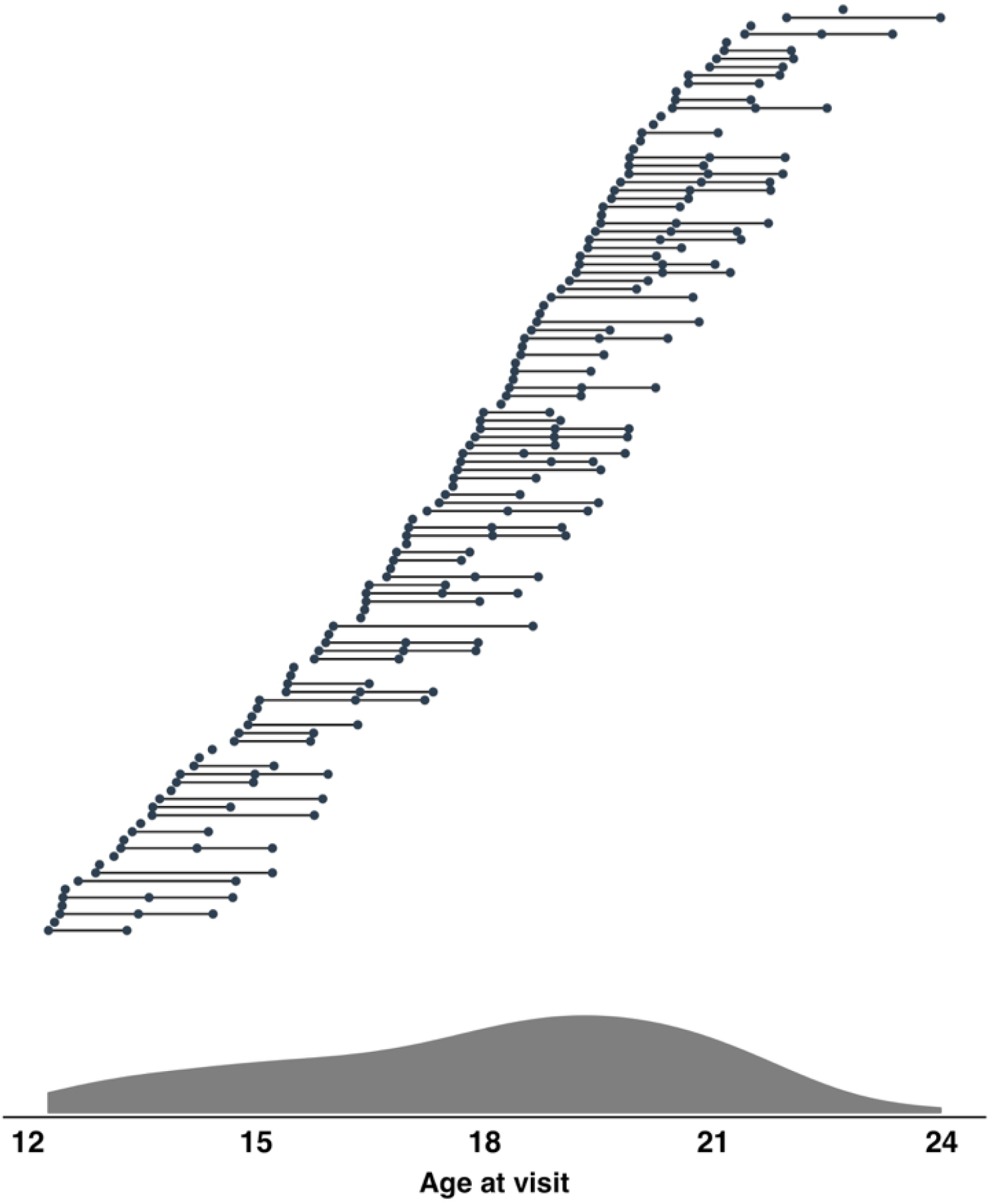
**Note:** Age distribution of sample. Primary behavioral sample included 113 participants and a total of 220 sessions. Top) Longitudinal structure of the project, where horizontal lines connect subjects’ visits (filled circles). Bottom) Density plot of subject age across all visits.

### 2.2 Measures

#### 2.2.1. Substance Use

The primary focus of the current study was to characterize developmental trajectories of substance use risk factors. Therefore, in order to account for inhibitory control deficits that may be associated with problematic substance use (Hardin & Ernst, 2009), problematic substance use was examined as a covariate in all analyses. Problematic substance use was defined at baseline using an established scoring protocol from the Customary Drinking and Drug Use Record (CDDR; Brown et al., 1998), where substance use was coded categorically and indicated whether participants exceeded age-specific National Institute on Alcohol Abuse and Alcoholism (NIAAA) guidelines for risky drinking (Exceeds Threshold Drinking; ETD; n = 29). This variable was selected for primary analysis to match our work with this sample using baseline and one-year follow-up data (Tervo-Clemmens et al., 2017). Furthermore, problematic substance use beyond this baseline characterization was uncommon in the final behavioral sample (weekly binge drinking: n = 17, weekly marijuana use n = 14; see **Supplementary Methods** for details), reflecting the isolation of substance use risk and relatively minimal levels of problematic substance use at this stage of the Pittsburgh NCANDA sample.

#### 2.2.2. Substance Use Risk Factors

Constructs indexing substance use risk in the current project, externalizing and internalizing psychopathology and family history of substance use disorder, were defined with regard to the original design of NCANDA (see Brown et al., 2015). Externalizing (EXT) and internalizing (INT) psychopathology were assessed using the Achenbach System of Empirically Based Assessments (ASEBA; Achenbach, 2009). Participants younger than 18 completed the Youth Self-Report (YSR), while participants older than 18 completed the Adult-Self-Report after the age of 18 (ASR). Critically however, these scales are designed to measure the same latent externalizing and internalizing structures, but with developmentally-appropriate questions. Age and gender adjusted t-scores were used as continuous measures for both EXT and INT. The ASEBA measures were collected at each visit. Family history of substance use disorder was assessed using the Family History Assessment Module (Rice et al., 1995), and was defined at baseline as having at least one biological parent or grandparent with substance use disorder symptoms (FH; *n* = 22). See Table 2 for correlation amongst substance use risk factors and other study variables.

**Table 2.** Correlations among risk and sociodemographic factors of interest

|  | EXT | INT | FH | ETD | Age | SES | GA | Gender |
| --- | --- | --- | --- | --- | --- | --- | --- | --- |
| EXT | — | Pearson | Polyserial | Polyserial | Pearson | Pearson | Pearson | Polyserial |
| INT | **0.517\*\*\*** | — | Polyserial | Polyserial | Pearson | Pearson | Pearson | Polyserial |
| FH | 0.184 | 0.134 | — | Polychoric | Polyserial | Polyserial | Polyserial | Polychoric |
| ETD | **0.298\*\*** | 0.012 | 0.181 | — | Polyserial | Polyserial | Polyserial | Polyserial |
| Age | 0.098 | 0.066 | -0.005 | **0.520\*\*\*** | — | Pearson | Pearson | Polyserial |
| SES | **-0.178\*** | -0.118 | -0.142 | 0.067 | 0.080 | — | Pearson | Polyserial |
| GA | 0.065 | -0.027 | 0.078 | 0.192 | **0.427\*\*\*** | **0.408\*\*\*** | — | Polyserial |
| Gender | 0.007 | -0.029 | **-0.338\*** | 0.021 | 0.129 | -0.035 | -0.044 | — |

**Note.** \* $p < 0.05$ , \*\* $p < 0.01$ , \*\*\* $p < 0.001$ . Categorical variables (FH and ETD) were coded as 1 for meeting risk criteria. Gender was coded with females as 1. Correlation coefficients (person: continuous-continuous associations; polyserial: continuous-categorical associations; polychoric: categorical-categorical) were generated from the polycor R package (Fox et al., 2010). Significance was determined using Pearson correlation for continuous-continuous associations, Welch's t-test for continuous-categorical associations, and chi-square testing for categorical-categorical associations.

#### 2.2.3. Generalized Cognitive Ability and Sociodemographic Factors

A composite measure of cognitive function, generalized cognitive ability accuracy (GA), was assessed with the Penn Computerized Neurocognitive Battery (Gur et al., 2010) and “paper and pencil” neuropsychological tests (Sullivan et al., 2016). GA was used as a covariate (see **2.5** for covariate procedures) in analysis of inhibitory control performance and brain activation in order to provide broader context of any developmental patterns of substance use risk factors associated with antisaccade performance. Specifically, since the GA measure evaluates several cognitive domains, this score allowed us to examine whether the developmental associations of substance use risk factors with antisaccade performance were specific to inhibitory control. Similarly, gender and Socioeconomic Status (SES; Hollingshead, 1975), which was measured as a composite of parental education and income, were also examined as covariates to provide broader sociodemographic context of substance use vulnerability (cf., Tarter et al., 2003). GA and SES information were defined at from baseline visit data. Sessions missing GA (n = 2) and SES (n = 14) data were excluded from analyses examining these variables.

### 2.3. Rewarded Antisaccade Task

Participants completed a rewarded antisaccade task during fMRI acquisition initially described by Geier et al (2010; Figure 2) and reported in prior work from our group on an earlier portion of this sample (Tervo-Clemmens et al., 2017). Briefly, each trial consisted of cue, preparation, and response epochs lasting 1.5s each. During the cue epoch, participants viewed a white fixation cross surrounded by a ring of green “$” symbols, indicating that a trial would be rewarded if a correct antisaccade was performed, or a ring of blue “#”, indicating a neutral trial in which no money was at stake. The preparation epoch was indicated by a red fixation cross and prompted participants to prepare for the response epoch, during which participants were to make a saccade in the mirror location of a yellow dot that appeared in unpredictable locations along the horizontal meridian at 1 of 6 eccentricities (±3, 6, and 9 degrees visual angle, relative to fixation). In addition to these “full trials”, additional partial trials, which consisted of either solely the cue epoch or the cue and preparation epochs but not the response epoch, were presented in order to aid in the estimation of hemodynamic responses evoked during each epoch (Ollinger, Corbetta, & Shulman, 2001; see **2.6.4**). Participants completed four neuroimaging runs, each consisting of 28 full trials (14 neutral, 14 reward) and 12 partial trials (3 of each partial trial by reward type). Prior to starting the task, participants were told that they could earn up to $25.00 but were not informed of the exact amount of monetary rewards for each correct trial. This was done in order to prevent participants from tallying their earnings, potentially engaging non antisaccade-specific brain systems (e.g., working memory).

**Figure 2.**
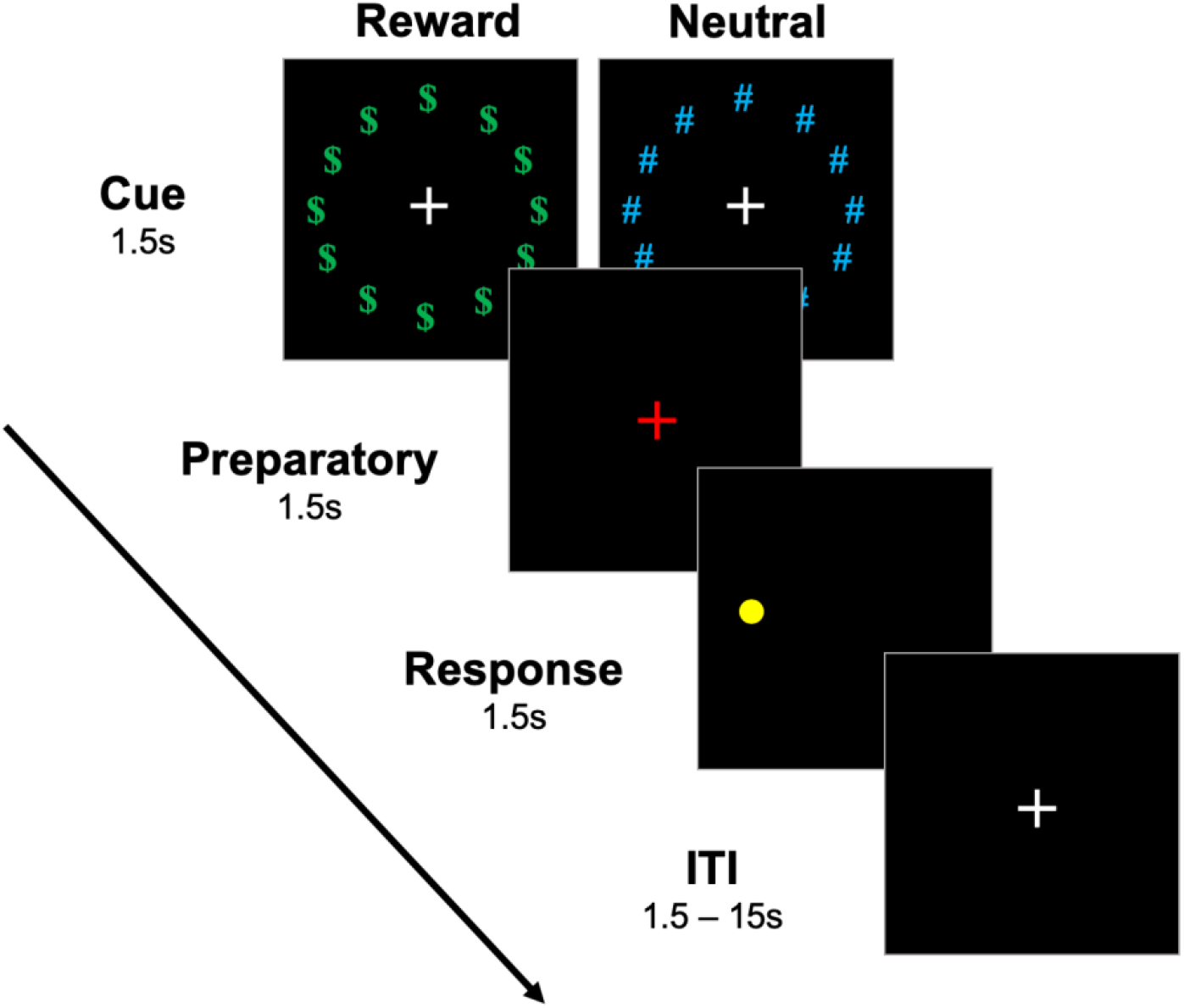
**Note.** Rewarded antisaccade task (Geier et al., 2009)

### 2.4. Eye Movement Measurement and Scoring

Antisaccade task stimuli were presented using E-Prime on a flat screen positioned behind the MRI scanner, which was visible to the participant through a mirror mounted on the MRI head coil. Eye movements were tracked using a long-range optics eye-tracking system (Applied Science Laboratories, Bedford, MA) that recorded eye positions through corneal reflections. At the beginning of each experimental session, a 9-point calibration was conducted to ensure accurate characterization of saccades for each participant.

Antisaccade scoring followed the same procedure as prior work from our group (Paulsen, et al., 2015, Tervo-Clemmens et al., 2017). A correct antisaccade was defined by an eye movement with a velocity of at least 30 degrees/s and 2.5 degree visual angle from central fixation to the mirror location of the peripheral target (Gitelman, 2002). Antisaccade errors were defined as those with an initial saccade toward the target that extended at least 2.5° visual angle (i.e. prosaccade). An error corrected trial was specified by an initial saccade toward the target and extended at least 2.5° visual angle from central fixation followed by a correct antisaccade to the mirror location. In the current project, both antisaccade errors and error corrected antisaccades were considered errors in both behavioral and neuroimaging analyses. Trials in which the participant failed to make any saccade or had poor eye tracking were considered dropped trials and were excluded from all analyses. Testing sessions (subject at visit) were excluded from both behavioral and neuroimaging analyses if the proportion of dropped trials exceeded 30% and/or if 50% of total trials were dropped or missing due to incomplete acquisition. The overall proportion of dropped trials was low across visits (10.6% at baseline, 11.3% at the second visit, and 10.9% at the third visit) and was not associated with substance use risk factors (**Supplementary Table 1**).

### 2.5. Antisaccade Behavioral Performance

Behavioral analyses were performed in R 3.1.2 (Rstudio Team, 2016). Generalized mixed-effects regression models with a logit link function (lme4 package in R, Bates et al., 2015) were used to predict antisaccade correct response rate across trial-wise data. This logit link function was appropriate considering the binomial structure of the trial-level data (correct vs. incorrect). Critically, the use of generalized mixed effect models with trial-wise data reduces ceiling effects in mean accuracy distributions (Dixon, 2008). Random intercepts were estimated for each subject. Reward conditions (reward, neutral), age, visit, ETD, and substance use risk factors (EXT, INT, FH) were entered as fixed effects in a series of models (see below). Antisaccade latencies were modeled similarly but using linear mixed effects models and only included correct trials. Significance values were obtained through the Car package in R (Fox et al., 2011). Potential leverage points were examined on session-mean performance metrics using cook’s distance with a threshold of 1. Only one session (subject at visit) exceeded this threshold and the pattern of significant results was unchanged when excluding this session. Consequently, this session was retained in the final analyses.

To examine main effects, the association between antisaccade performance and each risk factor was first modelled separately. Next, given the interest in studying developmental trajectories of inhibitory control among substance use risk factors, each risk factors’ interaction with age was examined. Subsequently, in order to address the specificity of results, all other risk factors were included as covariates. Lastly, to account for sociodemographic effects and general cognitive differences, GA and SES were included as covariates in both main effect and interaction models.

### 2.6. fMRI

#### 2.6.1. Acquisition

Neuroimaging data were collected at the Magnetic Resonance Research Center at the University of Pittsburgh on a 3.0-T Siemens Magnetom TIM Trio. A magnetization prepared rapid acquisition gradient-echo (MP-RAGE) pulse sequence with 160 slices (1.2 x 0.938 x 0.938) was used to create structural images for functional registration and conversion into a standardized template. Blood-oxygen-level-dependent (BOLD) data were acquired using an axially acquired echo-planar imaging (EPI) sequence (field of view = 200mm, 64 x 64 matrix, 3.125 x 3.125 x 3.200 mm anisotropic voxels, 29 slices, TR = 1.5s, TE = 28 ms, flip angle = 73°).

#### 2.6.2. Preprocessing

Preprocessing of fMRI data followed the same procedures as other recent task-based fMRI from our group (Paulsen, et al., 2015; Simmonds et al., 2018; Tervo-Clemmens et al., 2017). This included the removal of motion and noise artifacts (Analysis of Functional Neuroimages, AFNI; 3dDespike; Cox, 1996), non-linear registration of functional data to a standardized structural brain (3 mm, MNI-152 template; 2009c), slice timing and motion correction (mcflirt; Jenkinson et al., 2002). Data was then smoothed with FWHM of 5mm (SUSAN; Smith & Brady, 1997) and a 0.00625 Hz high-pass filter was applied. Finally, data was scaled by 10,000 of the global median. Sessions were excluded if session-wise mean Euclidean norm head motion exceeded 0.9 mm (n = 5), there was poor EPI coverage across runs upon visual inspection (n = 3), or due to technical error (n = 8).

#### 2.6.3. Trial-level fMRI Analysis

Consistent with prior work using a subset of this sample (Tervo-Clemmens et al., 2017), we estimated BOLD responses at both the trial and epoch level. To estimate average trial-level BOLD responses, a general linear model estimating BOLD activation was generated using AFNI’s 3dDeconvolve tool. Trial types (reward, neutral) were modelled separately with a 4,500 ms boxcar convolved with a gamma function (AFNI’s block 4). Individual regressors for correct, error corrected, and dropped trials were modeled for each trial condition. We note that incorrect (non error-corrected: **see 2.4**) antisaccades were rare in the sample overall (less than 1% of total trials) and did not occur in all subjects (69/104). Therefore, in order to ensure consistency of modeling across all subjects, these trials were modeled with the dropped trials. Partial trials were modelled similarly but with a 1,500 ms boxcar for cue partial trials and 3,000 ms boxcar for cue and preparatory partial trials. Six rigid-body head motion parameters, their first order derivatives, as well as run-wise 0 through 3rd order polynomials were included as nuisance regressors. TRs were censored if Euclidean norm head motion exceeded 0.9 mm.

#### 2.6.4. Epoch-level fMRI Analysis

Previous research has shown that BOLD activation varies amongst the epochs of this rewarded antisaccade task as participants assess reward contingencies, prepare a response, and execute an antisaccade response (Geier et al., 2010). Therefore, we also estimated the BOLD response within individual epochs in a second model. Each epoch (cue, preparation, and response) was modeled as a separate regressor with a boxcar (1.5sec duration, scaled to an amplitude of 1; AFNI’s block 4) and convolved with a gamma function. Correct, error corrected, and dropped/incorrect trials were modeled separately for each trial condition (Reward, Neutral) and each epoch. Partial trials were included as examples of each epoch, in order to aid in the estimation of epoch-specific BOLD response (Ollinger, Corbetta, & Shulman 2001). Nuisance regressors and motion censoring methods followed the same procedure as the trial-level deconvolution.

#### 2.6.5. Voxelwise Analysis

We examined main effects of substance use risk factors and their interactions with age in whole-brain voxelwise analyses predicting BOLD activity during correct antisaccade trials (AFNI’s 3dlme; Chen et al., 2013). All fMRI analyses covaried for session-wise mean Euclidean norm head motion, visit number, and trial type (reward, neutral). Each model was tested on both trial-level and individual epoch-level BOLD parameter estimates. Voxelwise testing was limited to voxels with at least a 50% probability of being gray matter in the MNI-152 template and full EPI coverage across the sample. Multiple comparison correction was conducted using a combination of cluster size and voxel significance. Parameters for thresholding were determined using AFNI’s 3dclustim program (acf option), which determines cluster size threshold through Monte Carlo simulations based on averaged spatial autocorrelation parameters. Autocorrelation parameters were estimated from residuals of the subject-level deconvolution models with AFNI’s 3dfwhmx tool. This analysis specified that at least 20 contiguous voxels initially thresholded at p=.005 were necessary to achieve cluster-level corrected alphas of less than .05 for both trial-level and epoch-level analyses.

## 3. Results

### 3.1. Antisaccade Behavioral Performance

#### 3.1.1. Reward and age effects

The average antisaccade correct response rate (accuracy) was 78.85% (SD = 15.69 %) and the average latency was 438.53 ms (SD = 57.5 ms). Both accuracy (z = 4.98, *X2(1)* = 24.78, p < 0.001) and latency (t =-3.66, *X2(1)* = 13.36, p < 0.001) robustly improved with age (higher accuracies and faster latencies with increasing age). Consistent with prior work using this task (Geier et al., 2010) and analyses within a subset of this sample (Tervo-Clemmens et al., 2017), rewarded trials were associated with greater correct response rates (z = 9.23, *X2(1)* = 85.01, p < 0.001) and shorter latencies (t =-7.60, *X2(1)* = 57.37, p < 0.001). Given that our prior work with a subset of this sample (Tervo-Clemmens et al., 2017) and the current project revealed non-significant interactions between reward conditions and substance use risk factors (**Supplementary Table 2**), additional higher-order interactions with trial type (reward, neutral), age, and risk-factors, were not explored. However, trial type was treated as a covariate in all behavioral and neuroimaging analyses.

#### 3.1.2. Main effects of substance use risk factors

##### Accuracy

When considering main effects across the sample, both EXT (*z* = −2.05, *X2(1)* = 4.21, *p* = 0.04) and INT (*z* =-3.24, *X2(1)* =10.51, *p* = 0.001) negatively predicted AS correct accuracy, suggesting that higher levels of these risk factors are associated with poorer antisaccade performance. Only INT (*z* =-2.16, *X2(1)* = 4.68, *p* = 0.02) continued to negatively predict AS accuracy when all risk factors were considered in a joint model. Both EXT (*z* = −2.15, *X2(1)* = 6.31, *p* = 0.01) and INT (*z* = −3.58, *X2(1)* = 12.82, *p* < 0.001) continued to be negatively associated with AS accuracy when covarying for GA and SES. FH and ETD did not predict AS accuracy (Table 3).

**Table 3.** Main effects and age interactions predicting antisaccade performance

|  |  | EXT | INT | FH | ETD | Age | GA | SES | Gender |
| --- | --- | --- | --- | --- | --- | --- | --- | --- | --- |
| Accuracy<br>(z) | Main effect | **-2.05\*<sub>b</sub>** | **-3.24\*\*<sub>a,b</sub>** | -1.13 | 1.02 | **4.98\*\*\*<sub>a,b</sub>** | **2.94\*\*<sub>a,b</sub>** | 0.60 | -0.06 |
|  | Age interaction | **2.48\*<sub>a,b</sub>** | 1.35 | -1.88 | -0.38 | — | **-3.08\*\*<sub>a,b</sub>** | **-2.66\*\*<sub>a,b</sub>** | -0.05 |
| Latency<br>(t) | Main effect | -1.20 | **-3.01\*\*<sub>a,b</sub>** | -1.10 | 0.21 | **-3.66\*\*\*<sub>a,b</sub>** | 0.88 | 0.43 | **3.82\*\*<sub>a,b</sub>** |
|  | Age interaction | **3.63\*\*\*<sub>a,b</sub>** | 0.47 | -0.10 | 0.77 | — | 1.39 | -.43 | -1.0 |

**Note.** \* $p < 0.05$ , \*\* $p < 0.01$ , \*\*\* $p < 0.001$ . Displayed test statistics are from models with the specific factor, age, visit, and trial condition (Reward, Neutral). <sub>a</sub>Significant relationships when covarying for all risk factors. <sub>b</sub>Significant relationships ( $p < .05$ ) when covarying for generalized cognitive ability (GA) and socioeconomic status (SES). Significant effects are bolded.

##### Latency of correct trials

Greater INT scores (*t* =-3.01, *X2(1)* = 9.04, *p* = 0.003) were associated with shorter latencies and remained a significant predictor of latency when considering all other risk factors (*t* = −2.22, *X2(1)* = 4.95, *p* = 0.006). Further, INT (*t* = −3.62, *X2(1)* = 13.14, *p <* 0.001) continued to predict faster latencies when controlling for GA and SES. EXT, FH, and ETD were not associated with main effects of latency (Table 3). As in the accuracy analysis, no other risk factors had significant interactions with age to predict latency.

#### 3.1.3. Risk factor interactions with age

##### Accuracy

EXT significantly moderated age-related improvements in AS accuracy (*z* = 2.48, *X2(1)* = 6.15, *p* = 0.013, Figure 3a). The Johnson-Neyman technique, which specifies intervals of significance within continuous by continuous interaction terms (Johnson & Fay, 1950), indicated that greater EXT scores predicted poorer AS accuracy in early adolescence until normalizing to the rest of the sample at approximately 17 years old (Figure 3a). The interaction between EXT and age remained significant when accounting for INT, FH and ETD (*z* = 2.64, *X2(1)* = 6.99, *p* = 0.008), as well as when covarying for GA and SES (*t* = 2.00, *X2(1)* = 3.99, *p* = 0.046). No other risk factors had significant interactions with age.

**Figure 3.**
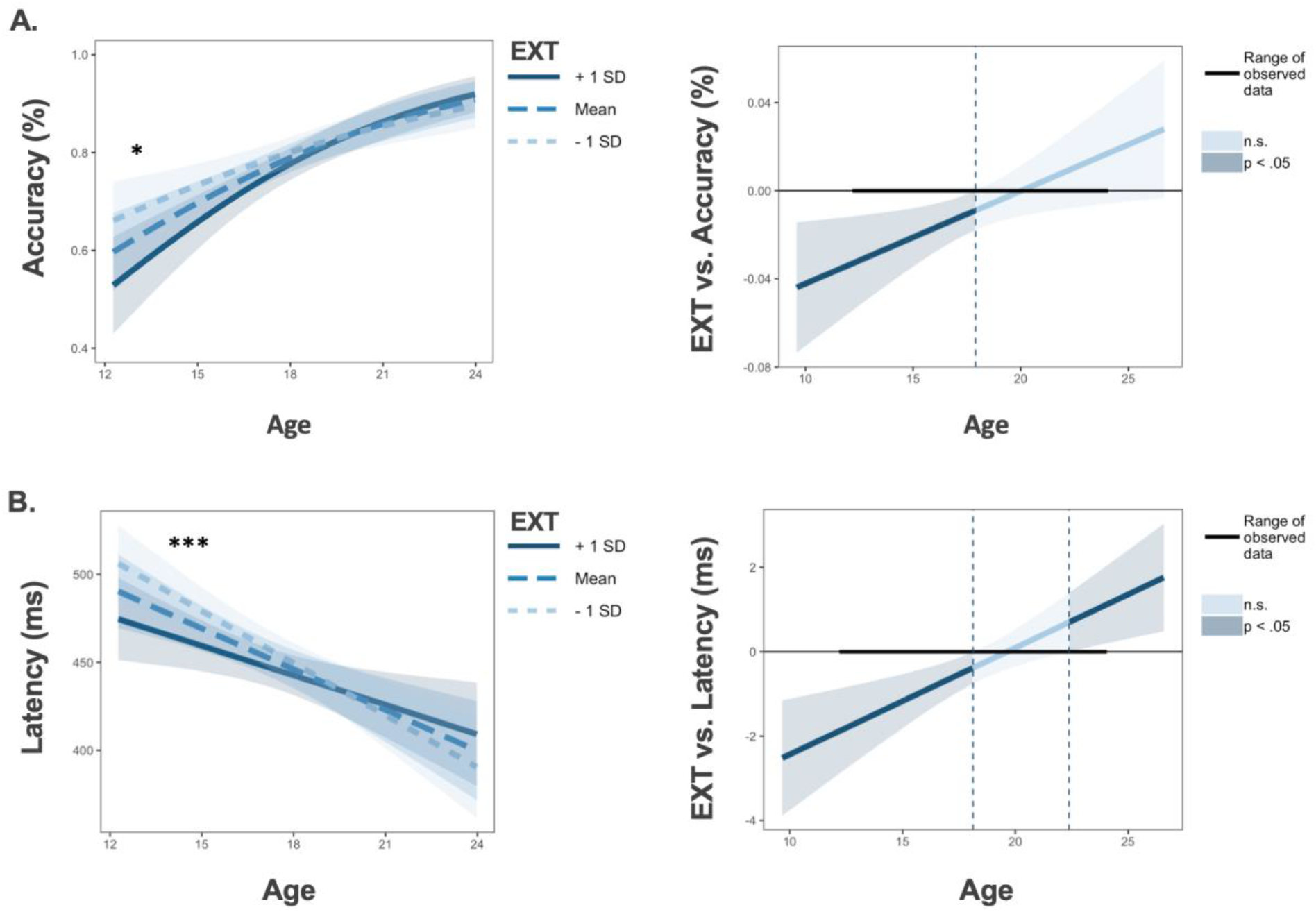
**Note:** *p < 0.05, **p < 0.01, ***p < 0.001. Individual differences in externalizing psychopathology (EXT) moderate age-related improvements in antisaccade accuracy (Panel A, left) and latency (Panel B, left). For Johnson-Neyman plots (generated from interactions R package; Long et al., 2019), darker blue colors reflect significant effects of EXT (p < 0.05) predicting antisaccade performance during the indicated age ranges (Panel A Panel B right). Shaded regions represent 95% confidence intervals.

##### Latency of correct trials

EXT also moderated age-related change in latency, where greater EXT scores were associated with reduced latency in early adolescence (*t* = 3.63, *X2(1)* = 13.18, *p* < 0.001; Figure 3b). This relationship persisted when covarying for all other risk factors (*t* = 3.96, *X2(1)* = 15.69, *p* < 0.001) and when accounting for GA and SES (*t* = 2.84, *X2(1)* = 8.06, *p* = 0.004). As in the accuracy analysis, no other risk factors had significant interactions with age.

##### Speed-Accuracy Relationships in Externalizing

Given the evidence for high externalizing subjects to have lower accuracy and faster latencies in correct responses during early adolescence, we performed a secondary analysis to examine potential speed-accuracy tradeoffs as a function of externalizing and age. This analysis revealed a significant three-way interaction between latency, age, and externalizing when predicting correct response rate (*z* = 3.04, *X2(1)* = 9.27, *p* = 0.002). However, visualization of accuracy and latency as a function of age and externalizing score suggested that externalizing performance differences in early adolescence were not consistent with a speed-accuracy tradeoff, as the largest accuracy difference associated with externalizing were observed for longer latencies (see **Supplementary Figure 1.**).

#### 3.1.4. Sociodemographic relationships

Females had significantly faster latencies (*t* = 3.82, *X2(1)* = 14.55, *p* < 0.001). Greater GA scores were associated with improved accuracy (*z* = 2.94, *X2(1)* = 8.64, *p* = 0.003). Further, GA (*z* = −3.08, *X2(1)* = 9.40, *p* = 0.002; **Supplementary Figure 2a.**) and SES (*z* = −2.66, *X2(1)* = 7.09, *p* = 0.008; **Supplementary Figure 2b.**) interacted with age to predict antisaccade accuracy, with patterns that mirrored the interaction between EXT and age.

### 3.2. Functional Neuroimaging

#### 3.2.1. Head Motion

As has been well-described in previous developmental research, head motion was greater at younger ages (*t* =-2.74, *X2(1)* = 7.55, *p* < 0.001). No risk factors were associated with head motion or interacted with age to predict head motion. However, as described in the methods section, TRs during which Euclidean norm head motion distances exceeded 0.9 mm were excluded and all fMRI analyses covaried for average session-wise Euclidean norm head motion.

#### 3.2.2. Main effects of substance use risk factors

In the interest of studying developmentally-relevant brain-behavior relationships, only significant main effects and risk factor by age interactions from the behavioral analyses were followed-up in neuroimaging analyses. At the trial-level, greater EXT scores predicted decreased activation in the left middle frontal gyrus (Figure 4; **Supplementary Table 3**). Average activation within this cluster had a positive relationship with antisaccade accuracy (proportion of correct trials; *t* = 2.50, *X2(1)* = 6.23, *p* = 0.01), suggesting hypo-activation in this region was task-relevant to performance. EXT was also a negative predictor of BOLD activation in the right superior temporal gyrus (**Supplementary Table 3**). Among individual epochs, EXT was positively associated with BOLD activity within the thalamus during the preparatory epoch and had negative relationships with BOLD activation across frontoparietal regions during the response epoch (**Supplementary Table 3**).

**Figure 4.**
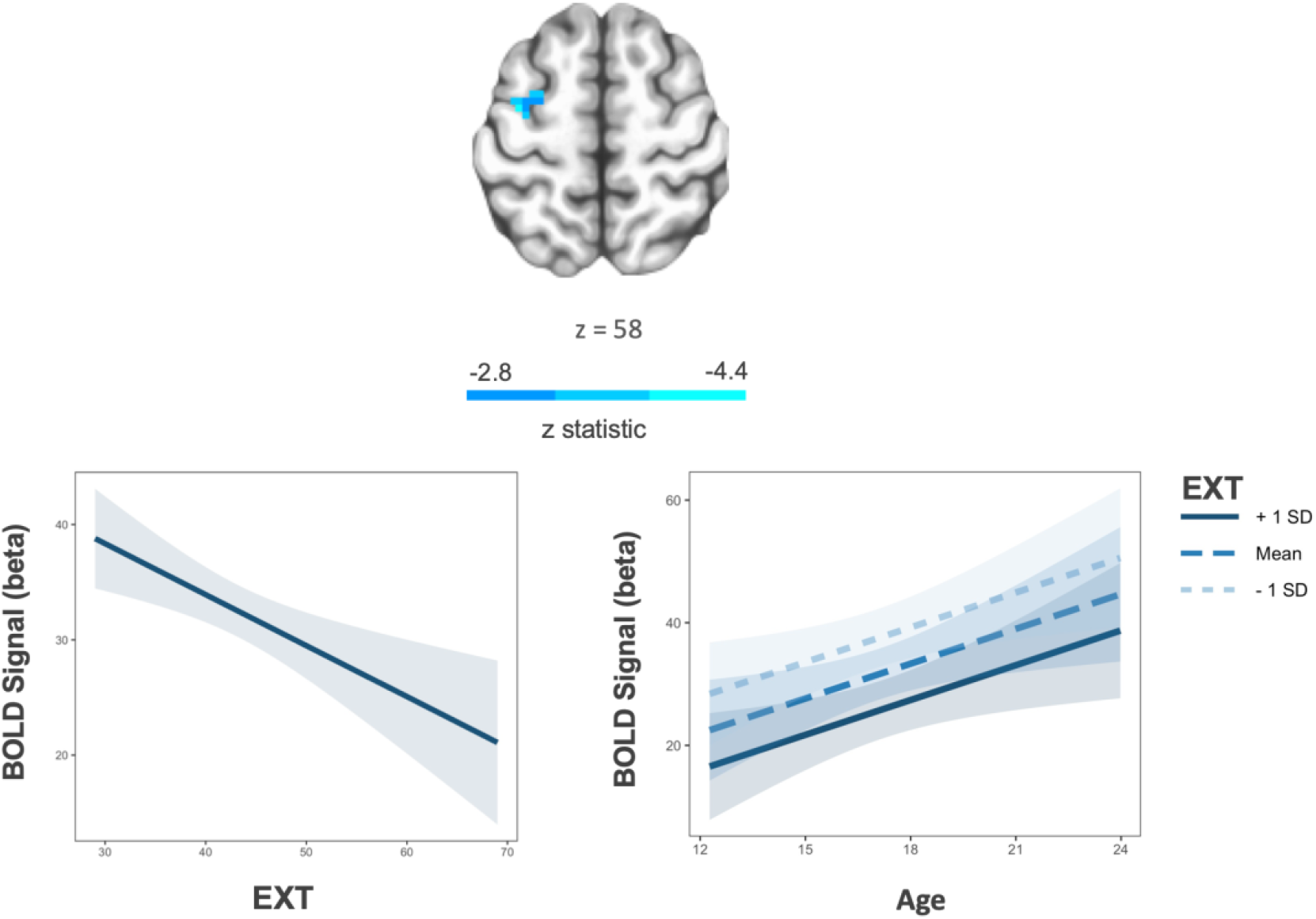
**Note:** Greater externalizing scores are associated with decreased trail-wise BOLD activation in the left middle frontal gyrus (left). This association did not significantly interact with age (right). Shaded regions represent 95% confidence intervals. Voxelwise threshold p < .005, number of contiguous voxels > 20, p < 0.05 corrected.

Specifically among trial-level data, greater INT scores were associated with decreased activity in frontoparietal regions (inferior parietal lobule, precuneus, middle frontal gyrus; **Supplementary Table 3**) and greater GA scores predicted increased activity in the left precuneus (**Supplementary Table 3**).

#### 3.2.3 Risk factor and sociodemographic interactions with age

At the trial-level, there was a significant GA and age interaction in the right middle frontal gyrus (Table 4; **Supplementary Figure 3**) such that greater GA scores predicted decreased activity in this region with increasing age. Average activation within this cluster was a positive predictor of antisaccade accuracy (*t* = 2.25, *X2(1)* = 5.04, *p* = 0.025). Further, SES interacted with age to predict BOLD activity in the inferior parietal lobule such that greater SES was associated with decreased activity in this region with increasing age (Table 4; **Supplementary Figure 3**).

**Table 4.** Age interactions of substance use risk and sociodemographic factors in BOLD activation

| Variable | Epoch | Region | BA | k | MNI |  |  | t |
| --- | --- | --- | --- | --- | --- | --- | --- | --- |
|  |  |  |  |  | x | y | z |  |
| EXT | Cue | L Posterior Parietal Cortex | 7 | 22 | -29 | -71 | 31 | 4.26 |
| GA | Trial-wise | R Middle frontal gyrus | 6 | 26 | 31 | -8 | 61 | -4.27 |
|  | Preparatory | R Precuneus | 31 | 20 | 13 | -59 | 22 | -4.14 |
|  |  | L Precuneus | 7 | 20 | -2 | -74 | 37 | -3.89 |
|  | Response | R Insula | 13 | 83 | 40 | -8 | 19 | 5.44 |
|  |  | L Precuneus | 31 | 20 | -20 | -62 | -25 | 4.03 |
| SES | Trial-wise | L Inferior Parietal Lobule | 40 | 24 | -59 | -41 | 25 | -4.37 |
|  | Cue | L Angular gyrus | 39 | 20 | -53 | -68 | 34 | 3.71 |
|  | Preparatory | L Inferior Parietal Lobule | 22 | 34 | -59 | -38 | 22 | -5.36 |

**Note.** BA, Brodmann Area; K, number of voxels in cluster. X, Y,Z, peak voxel coordinates in MNI space. t, mean cluster t-values from models predicting activation within clusters with a significant age interaction (voxelwise threshold $p < 0.005$ , number of contiguous voxels $\geq 20$ , $p = 0.05$ corrected). All voxelwise models covaried for trial type (reward, neutral), visit, and session-wise average head.

At the epoch-level, EXT moderated age-related BOLD activity within the left posterior parietal cortex (Table 4; Figure 5) in the cue epoch. Mean activation within this cluster did not predict antisaccade accuracy (proportion of correct trials; *t* = 0.46, *X2(1)* =0.21, *p* = 0.65) or latency (*t* = 1.24, *X2(1)* =15.40, *p* = 0.21). Mirroring similarities with EXT in behavioral data, GA and SES also interacted with age to predict BOLD activation within individual epochs. Specifically, greater GA scores were associated with decreased activation with increasing age in bilateral precuneus in the preparatory epoch, but were associated with increased activation with increasing age in the left precuneus and right insula during the response epoch (Table 4; **Supplementary Figures 4**). Greater SES scores were associated with increased activity in the left angular gyrus with increasing age during the cue epoch but decreased activity with increasing age in the left inferior parietal lobule during the preparatory epoch (Table 4; **Supplementary Figures 5**).

**Figure 5.**
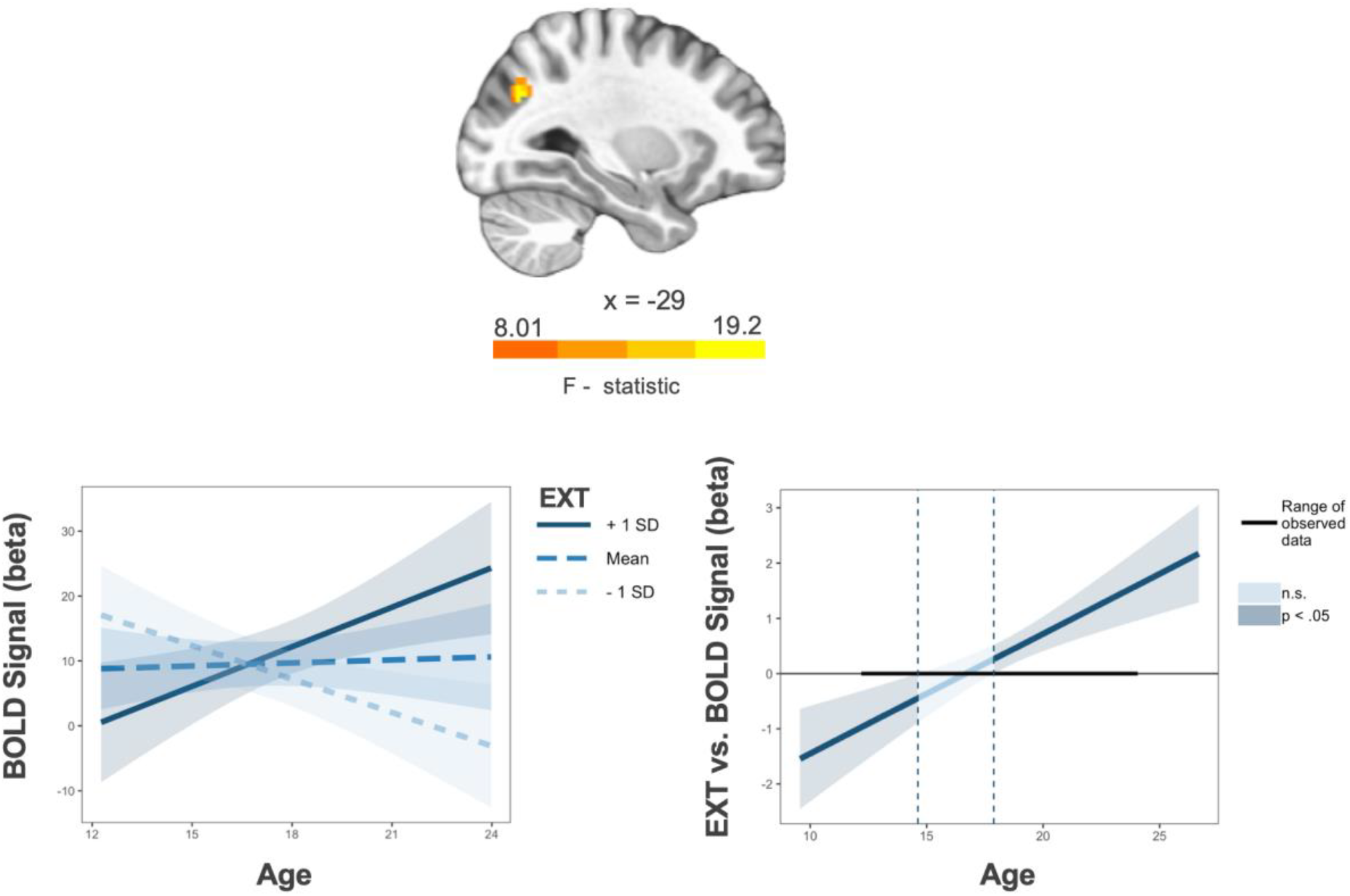
**Note:** Externalizing psychopathology (EXT) by age interaction in the left posterior parietal cortex during the cue epoch. Voxelwise threshold p < .005, number of contiguous voxels > 20, p < 0.05 corrected (left). For Johnson-Neyman plot of interaction, darker blue colors reflect significant main effects of EXT (p < 0.05) predicting BOLD signal during the indicated age ranges (right). Shaded regions represent 95% confidence intervals.

## 4. Discussion

The purpose of the current study was to examine the relationship between age-related trajectories of inhibitory control and substance use risk factors. Our findings indicate that higher levels of externalizing psychopathology are associated with developmental differences in inhibitory control, characterized by poorer accuracy and shorter latencies in early adolescence. Neuroimaging results revealed that externalizing psychopathology was associated with both developmentally-stable hypo-activation in the middle frontal gyrus and developmentally specific age-related increases in posterior parietal cortex activation during early adolescence.

### 4.1. Substance use risk factors and developmental improvements in inhibitory control

We found that greater externalizing scores were associated with poorer correct response rate in the antisaccade task. This result is consistent with our previous findings within a subset of the current sample (Tervo-Clemmens et al., 2017), and supports theories relating disinhibitory phenotypes of substance use risk with poor response inhibition (Iacono et al., 2008; Young et al., 2009). We further sought to investigate how the relationship between inhibitory control limitations and substance use risk changes developmentally. Consistent with previous work, antisaccade accuracy and latency improved throughout adolescent development (Ordaz et al., 2013). However, our findings indicate divergent developmental trajectories of inhibitory control with respect to the degree of substance use risk factors. Specifically, we show that greater levels of externalizing psychopathology were associated with poorer antisaccade accuracy and faster latencies in early adolescence. However, secondary analyses revealed a nuanced relationship between latency and accuracy in high externalizing youth during early adolescence, where the largest differences in correct response rate where observed at longer latencies. Thus, these findings do not support a speed accuracy trade-off model of impulsivity in which high impulsivity is associated with rapid and error-prone responses in cognitive tasks (Dickman and Meyer, 1988). Rather, they may be consistent with differences in an overall response strategy that differentially emphasizes speed or accuracy, depending on task demands. Our results further suggest such response differences are most evident in early adolescence, followed by a pattern of equifinality. However, despite poor response inhibition in early adolescence, high externalizing was associated with accelerated age-related improvements in accuracy to establish normative performance levels by late adolescence. Similarly, shorter latencies in early adolescence normalized by late adolescence. These developmental differences may reflect shifts in strategies to optimize performance throughout development and compensate for inhibitory control limitations in early adolescence, as high externalizing gain more top-down control of behavior. To this end, additional work may more directly test developmental changes in speed-accuracy tradeoffs and decision-making strategies in more explicit computational frameworks (e.g., drift diffusion modeling) in externalizing youth.

Although previous work implicates cognitive control deficits in youth with a family history of substance use disorder (Cservenka et al., 2015), this risk factor was not associated with differences in antisaccade performance in the current project. However, our analyses with this risk factor had limited statistical power due to few family history positive subjects within the sample. In contrast, and relatively unexpectedly, internalizing psychopathology was associated with low correct response rate and faster latencies across development. More clarity on family history and internalizing psychopathology within inhibitory control paradigms will require future work examining broad multidimensional high-risk profiles (see **4.3**). Nevertheless, externalizing was the only substance use risk factor that moderated age-related change in antisaccade performance, providing some support that externalizing may be uniquely associated with developmental differences in inhibitory control. To this end, the externalizing by age interaction remained significant when covarying other risk factors. Taken together, these results highlight early adolescence as a potential period of increased substance use vulnerability by way of developmental differences in response inhibition and disinhibitory phenotypes.

### 4.2. Substance use risk factors and brain systems supporting inhibitory control

Converging with previous research examining prospective prediction of substance use initiation and high-risk profiles, internalizing and externalizing substance use risk factors in the current project were associated with decreased BOLD activation across frontoparietal regions during response inhibition (Norman, et al., 2011; McNamee et al., 2008). Further, supporting previous work implicating poor top-down control in externalizing psychopathology (Beauchaine, et al., 2017), greater levels of externalizing were associated with decreased trial-level activation in the left middle frontal gyrus. Given that this finding occurs at the trial-level and activation in this region predicted antisaccade accuracy, poor inhibitory control performance associated with high externalizing may be marked by limitations in broad inhibitory control processes mediated by the middle frontal gyrus. Further, this relationship may represent a general trait-level feature of externalizing as it was stable across development. This relationship may also index a trait-level feature of general psychopathology as internalizing psychopathology was also associated with developmentally-stable decreased activation in frontoparietal regions. In contrast to common developmentally-stable effects, externalizing was uniquely associated with developmental differences in posterior parietal cortex activity during the cue epoch of the antisaccade task, where high externalizing was associated with increasing activation with age in this region. The developmental association of posterior parietal cortex activation with externalizing as a substance use risk factors is consistent with previous research showing that posterior parietal cortex activation differences are associated with adolescent substance use (Schweinsberg et al., 2008) and substance use vulnerability (Tervo-Clemmens et al., 2018a) and age of substance use initiation (Tervo-Clemmens et al., 2018b) during working memory tasks. Although the sign of activation may differ based on task demands, activation differences in the posterior parietal cortex may represent a common developmental neural correlate underlying substance use risk across cognitive tasks. Supporting this, the posterior parietal cortex is implicated in attentional control, a function supporting both working memory and inhibitory control (Corbetta & Schulman, 2002; Unsworth, 2004). Given that this finding occurs during the cue epoch of correct antisaccades, greater levels of externalizing may be associated with neurodevelopmental limitations in attentional control required when orienting to task demands.

Previous work characterizing normative functional development of the parietal cortex indicate this region does not exhibit age-related changes during the antisaccade task (Ordaz et al., 2013) but its function decreases with age to support working memory development (Simmonds et al., 2017). Diverging from these patterns, greater levels of externalizing were associated with increasing engagement of the posterior parietal cortex across development that may reflect a compensatory mechanism, not observed for low levels of externalizing and in more normative subjects (cf., Ordaz et al., 2013), that underlies the normalization of antisaccade performance in later adolescence and adulthood. However, in the absence of a significant association between activity in this region and antisaccade performance, the functional contribution of parietal cortex activity to developmental differences in inhibitory control performance associated with externalizing is unclear. Future work may use multiple tasks to examine general and specific activation patterns in the posterior parietal cortex that underlie the development of cognitive control in risk for substance use.

### 4.3. Common and specific neurocognitive profiles of substance use risk factors

In addition to characterizing development trajectories associated with substance use risk factors, we investigated developmental differences in broader sociodemographic factors, including generalized cognitive ability accuracy and socioeconomic status. Mirroring developmental differences associated with externalizing, lower levels of generalized cognitive ability and socioeconomic status were associated with decreased antisaccade correct response rates in early adolescence that normalized by late adolescence and early adulthood. These results indicate a potential common developmental pattern of equifinality across sociodemographic and psychopathology risk factors and support theories suggesting substance use risk may best be represented using dimensional constructs encompassing multiple facets of risk factors (e.g., neurobehavioral disinhibition; Tarter et al., 2003). However, our results also suggest that each risk factor may be associated with independent developmental processes underlying inhibitory control maturation. Supporting this, differences in age-related improvements in antisaccade *latency* was specific to externalizing. Further, generalized cognitive ability, socioeconomic status, and externalizing were associated with developmental differences in non-overlapping brain regions. Specifically, greater levels of generalized cognitive ability and socioeconomic status were associated with BOLD activity that decreased with age in the middle frontal gyrus and inferior parietal lobules, respectively. Thus, our data indicate potential compensatory mechanisms may be differentially relevant to each risk factor. Future research may further evaluate individual differences in neurocognitive mechanisms underlying developmental patterns of equifinality in inhibitory control common across substance use risk and sociodemographic factors.

### 4.4 Limitations

The present study is characterized by a number of strengths, including a relatively large longitudinal neuroimaging sample and use of continuous measures of substance use risk factors. This approach allowed us to identify the relative magnitude of substance use risk factors and distinct age ranges in which inhibitory control deficits are most evident. Such a characterization of individual differences in inhibitory control across development and among multiple risk factors highlights the importance of studying dimensional constructs of psychopathology (Research Domain Criteria; Morris & Cuthbert, 2012). However, in the current sample, a relatively small proportion of the sample exceeded the described threshold of alcohol use at baseline, and few participants initiated problematic substance use at follow-up visits. This limited our ability to examine how problematic substance use initiation and escalation may perturb normative development of inhibitory control. Similarly, relatively few participants met criteria for a family history of substance use disorder. Therefore, our study was less powered compared to previous studies that have characterized developmental differences in youth with a family history of substance use disorder (Hardee et al., 2014). With respect to previous work highlighting cognitive control deficits in youth with a family history of substance use disorder (Cservenka, 2016) and substance use initiation in adolescence (Hardin & Ernst, 2009), future research should consider broader sampling of youth with a family history of substance use disorder and greater longitudinal time points to better characterize developmental differences associated with these substance use risk factors and subsequent substance use escalation.

## 5. Conclusion

Our findings indicate that substance use risk factors are associated with developmental differences in inhibitory control. Specifically, externalizing psychopathology is associated with decreased correct response rate and shorter latencies in early adolescence that appears to resolve into adulthood, perhaps through compensatory mechanisms acquired during adolescence. Neuroimaging data reveal high externalizing is associated with developmentally-stable hypo-activation in prefrontal cortex and divergent development of posterior parietal cortex activation. Taken together, our data suggest early adolescence may be a unique period of substance use vulnerability via cognitive and phenotypic disinhibition.

## Acknowledgments

The authors acknowledge the participants and their families. AQ acknowledges Dr. A.W. Gilmore for thoughtful discussion and helpful comments on earlier drafts of the manuscript.

