## Supplementary Material for "Neurocognitive development of inhibitory control and substance use vulnerability"

### Supplementary Materials

**Supplementary Methods.** The NCANDA sample was designed, in part, to isolate *risk* for problematic substance use, with the majority of subjects recruited prior to significant substance use involvement (see Brown et al. 2015). Therefore, early time points in the longitudinal study are predominated by low levels of substance use and our primary analyses utilized a baseline assessment of problematic substance (Exceeds Threshold Drinking; ETD; n = 29). This approach had the benefit of matching our prior work examining baseline and one-year follow up visits with this sample (Tervo-Clemmens et al., 2017). However, the current project additionally utilized data from the second-year follow-up, which increases the likelihood that some subjects have transitioned into problematic substance use.

Among the subjects comprising the final behavioral sample for the current project (n=113), twenty subjects total transitioned into problematic substance use (weekly binge drinking) between either the baseline and one-year follow-up (n=13) or between the one- and two-year follow-up (n=7). However, only fifteen of these subjects met all data quality criteria to be included in behavioral analyses. To this end, within the small set of problematic substance use in this sample, regular (at least weekly) binge drinking (n=17) was not associated with antisaccade correct response rate (accuracy)( $z = -0.19$, $X_{2(1)} = 0.03$, $p = 0.85$) or latency ($t = -1.182$, $X_{2(1)} = 1.49$, $p = 0.24$). Regular marijuana use (at least weekly) was also fairly uncommon among the subjects in our final behavioral analyses (n=14) and was not significantly associated with antisaccade accuracy ($z = -1.21$, $X_{2(1)} = 1.46$, $p = 0.23$) or latency ($t = -1.816$, $X_{2(1)} = 3.30$, $p = 0.07$).

**Supplementary Table 1.** Main effects and age interactions predicting proportion of excluded antisaccade trials

| | EXT | INT | FH | ETD | Age | GA | SES | Gender |
|---|---|---|---|---|---|---|---|---|
| Main Effects | 0.10 | 1.07 | -1.41 | -0.04 | 0.48 | 0.30 | 0.54 | -0.37 |
| Age interaction | 0.01 | -0.04 | 0.71 | -0.71 | — | -0.54 | -1.08 | 0.16 |

**Note.** Displayed test statistics are t values from models predicting proportion of excluded trials. All models covaried for age and visit.

**Supplementary Table 2.** Reward interactions predicting antisaccade performance

| | EXT | INT | FH | ETD | Age | GA | SES | Gender |
|---|---|---|---|---|---|---|---|---|
| Accuracy (z) | -0.18 | -0.02 | -1.93 | 0.007 | -1.10 | 0.48 | 0.84 | 0.64 |
| Latency (t) | 1.25 | 0.37 | -1.76 | -0.67 | -2.09* | 0.06 | 1.29 | 0.16 |

**Note.** Displayed test statistics are t values from models predicting accuracy and latency. All models covaried for age and visit.

**Supplementary Table 3.** Main effect of substance use risk and sociodemographic risk factors on BOLD activation at the trial-level and within individual epochs

| | | | | | MNI | | | |
|---|---|---|---|---|---|---|---|---|
| | | | BA | k | x | y | z | t |
| EXT | Trial-wise | R Superior temporal Gyrus | 22 | 30 | 55 | -44 | 13 | -5.05 |
| | | L middle frontal gyrus | 6 | 33 | -35 | -8 | 58 | -5.01 |
| | Preparatory | R Anterior thalamus | — | 23 | 4 | -2 | 7 | 4.14 |
| | | R Posterior thalamus | — | 20 | 7 | -20 | 4 | 3.97 |
| | Response | L Mid-cingulate | 32 | 75 | -5 | -23 | 46 | -5.32 |
| | | R Superior frontal gyrus | 9 | 60 | 22 | 46 | 37 | -4.84 |
| | | R Temporal parietal junction | 40 | 57 | 58 | -50 | 19 | -5.46 |
| | | R superior temporal gyrus | 22 | 55 | 58 | -38 | 4 | -4.26 |
| | | L Precuneus | 7 | 34 | -11 | -74 | 37 | -4.30 |
| | | R Putamen | — | 33 | 16 | 10 | -2 | -4.47 |
| | | R SMA | 6 | 28 | 28 | -8 | 49 | -4.21 |
| | | R Medial PFC | 32 | 21 | 4 | 46 | 4 | -4.31 |
| | | R ACC | 24 | 21 | 4 | 25 | 25 | -4.27 |
| INT | Trial-wise | L inferior parietal lobule | 40 | 65 | -68 | -29 | 37 | -5.80 |
| | | L inferior parietal lobule | 19 | 36 | 28 | -80 | 34 | -4.39 |
| | | R inferior parietal lobule | 5 | 23 | 31 | -44 | 61 | -4.79 |
| | | R Precuneus | 40 | 20 | -41 | -41 | 46 | -5.25 |
| | | R Middle frontal gyrus | 7/19 | 59 | -26 | -74 | 40 | 5.26 |
| GA | Trial-wise | L Precuneus | 22 | 30 | 55 | -44 | 13 | -5.05 |

**Note.** BA, Brodmann Area; K, number of voxels in cluster. X, Y,Z, peak voxel coordinates in MNI space. t, mean cluster t-values from models predicting activation within clusters with a significant age interaction (voxelwise threshold $p < 0.005$, number of contiguous voxels ≥ 20, $p = 0.05$ corrected). All models covaried for trial type (reward, neutral), visit, and session-wise average head motion in voxelwise models.

**Supplementary Figure 1.**

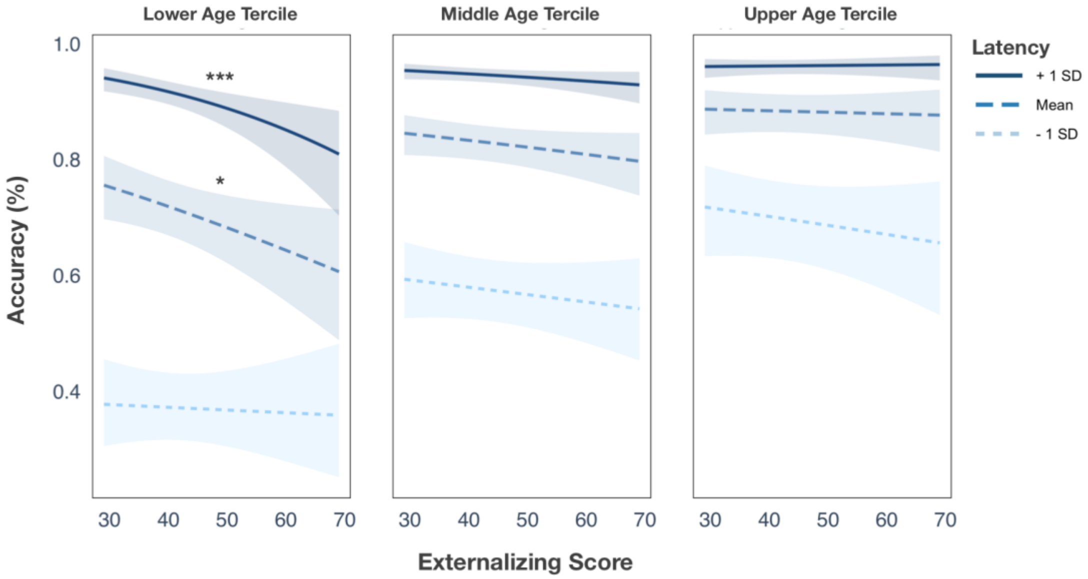


**Note.** Latency moderates the relationship between externalizing psychopathology and antisaccade accuracy in early adolescence. For visualization, age ranges are split into terciles represented in each panel (Median age in each tertiles are 15, 18.5, and 20.7 respectively). Shaded regions reflect 95% confidence intervals. Plots were generated from the *interactions* R package (Long et al., 2019).

**Supplementary Figure 2.**

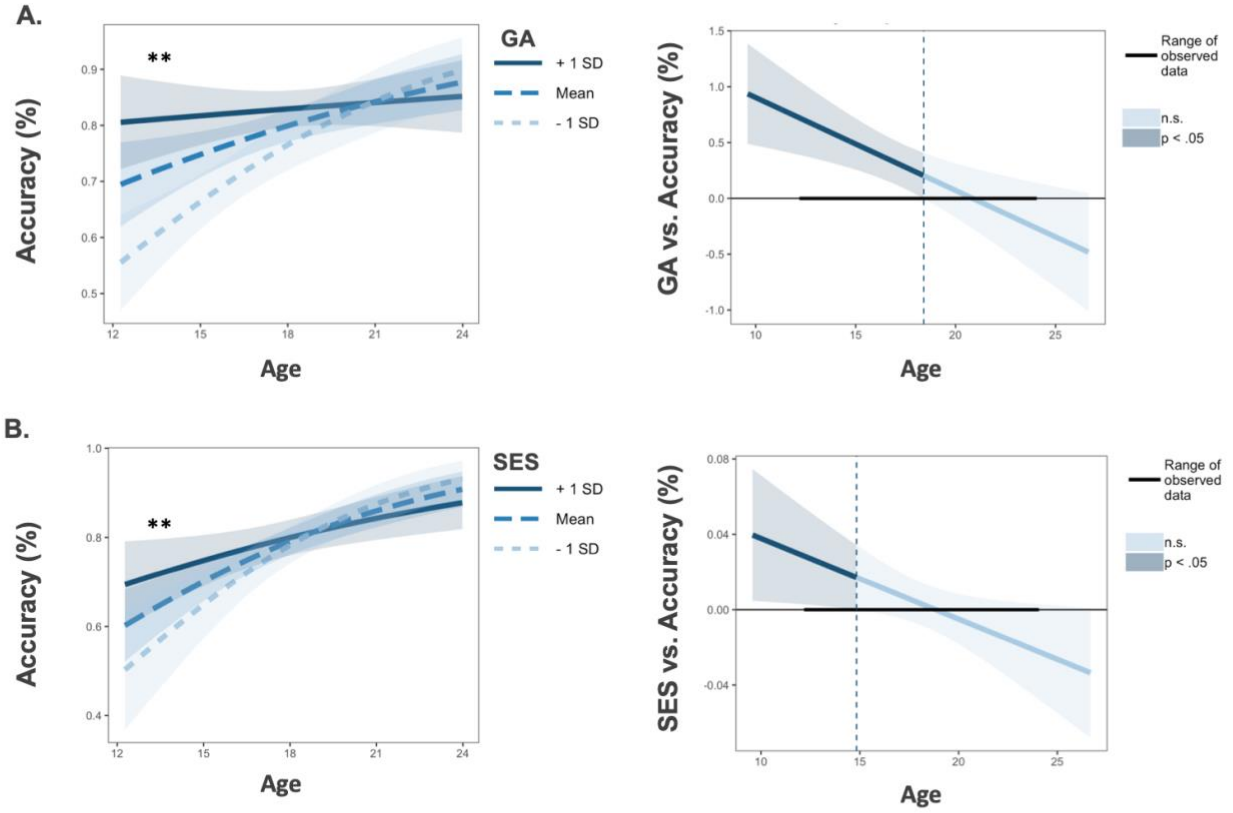


**Note.** Individual differences in generalized cognitive ability (GA, Panel A) and socioeconomic status (SES, Panel B) moderates age-related improvements in antisaccade accuracy. For Johnson-Neyman plots (right side), darker blue colors reflect significant main effects of GA or SES ($p < 0.05$) predicting antisaccade accuracy during the indicated age ranges. Shaded regions represent 95% confidence intervals.

**Supplementary Figure 3.**

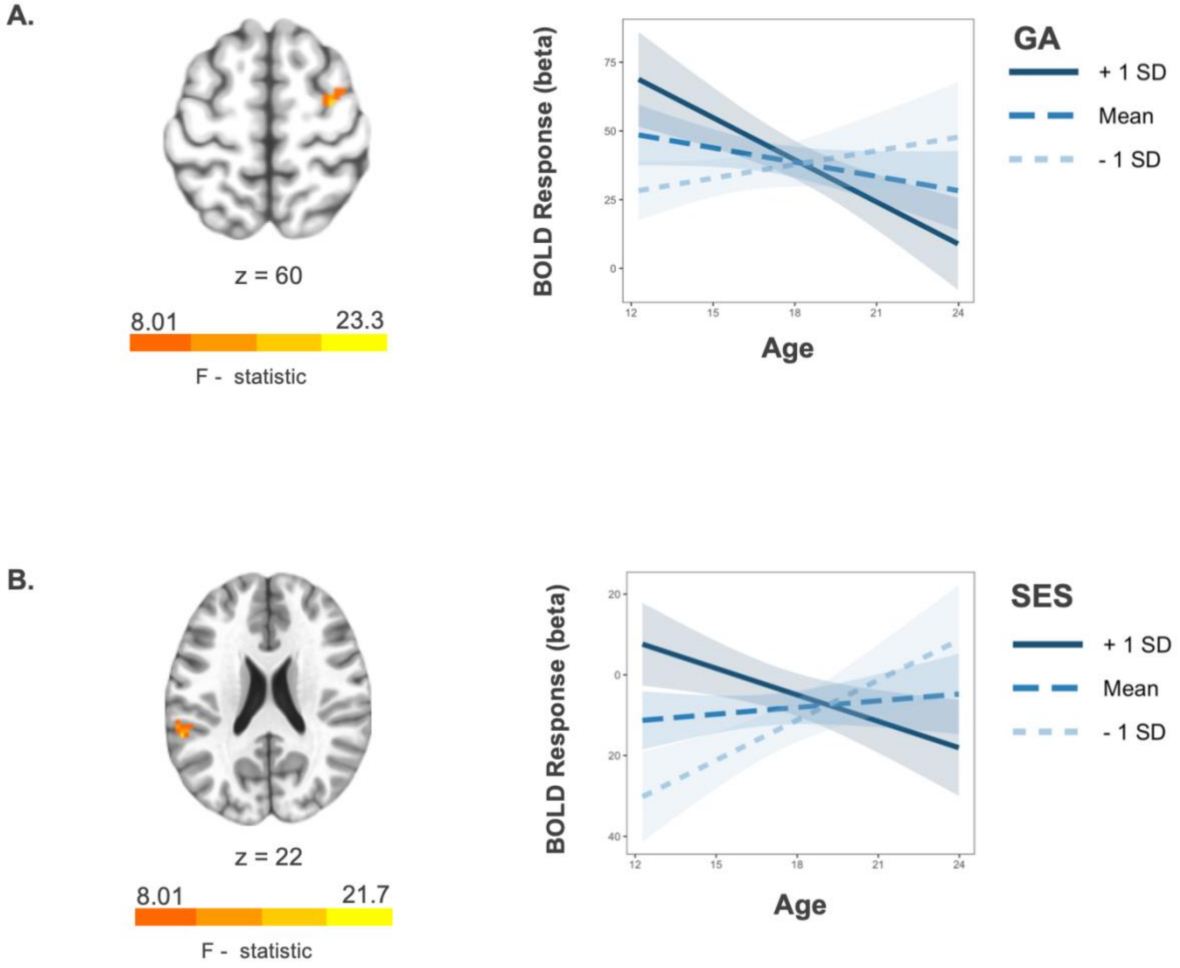


**Note.** Generalized cognitive ability (GA) by age interaction in the right middle frontal gyrus at the trial-level (Panel A). SES by age interaction in the left insula at the trial-level (Panel B). Voxelwise threshold $p < .005$, number of contiguous voxels > 20, $p < 0.05$ corrected.

**Supplementary Figure 4.**

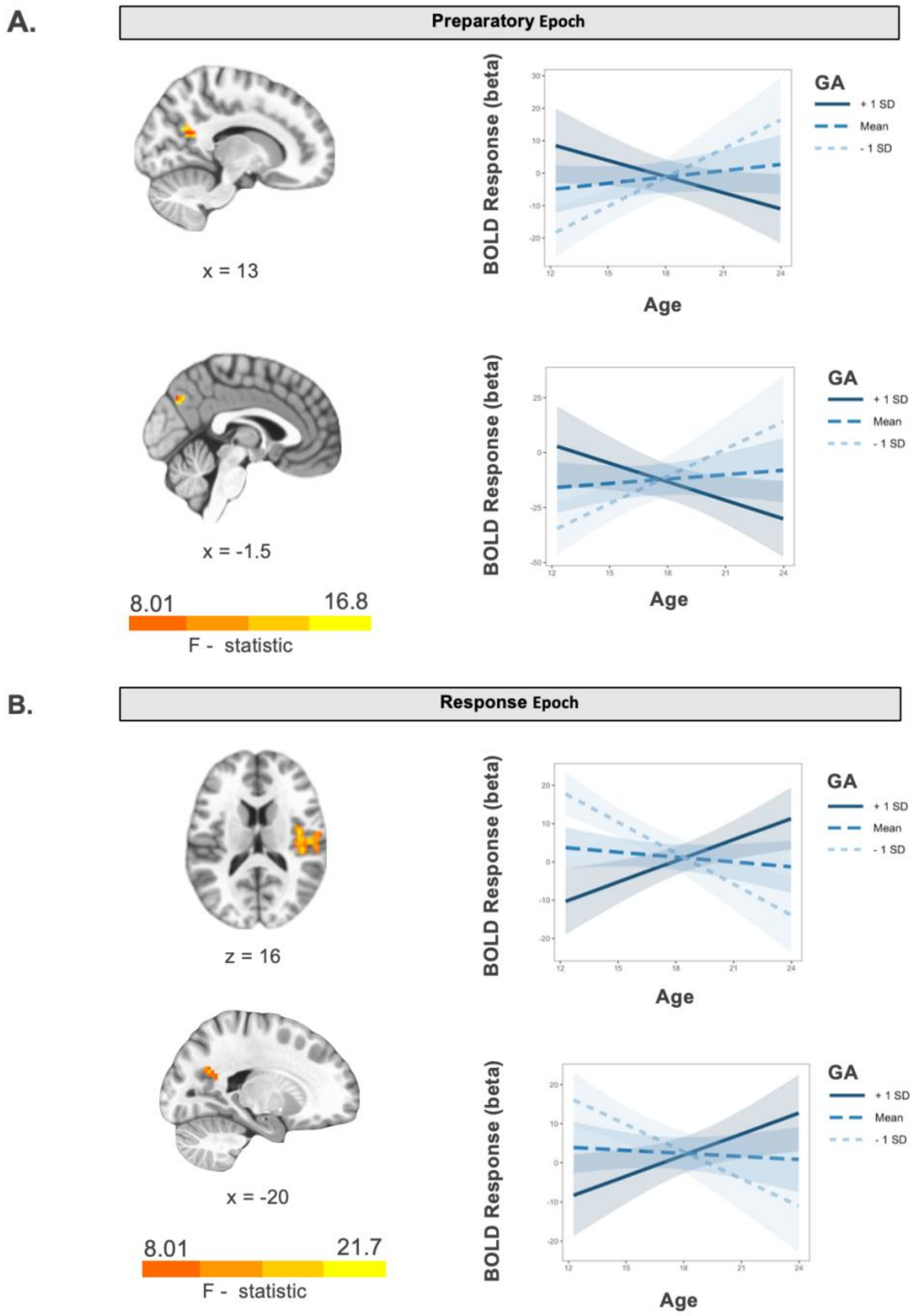


**Note.** Generalized cognitive ability (GA) by age interaction in bilateral precuneus during the preparatory epoch (Panel A). Generalized cognitive ability (GA) by age interaction in the right insula and left precuneus during the response epoch (Panel B). Voxelwise threshold $p < .005$, number of contiguous voxels > 20, $p < 0.05$ corrected.

**Supplementary Figure 5.**

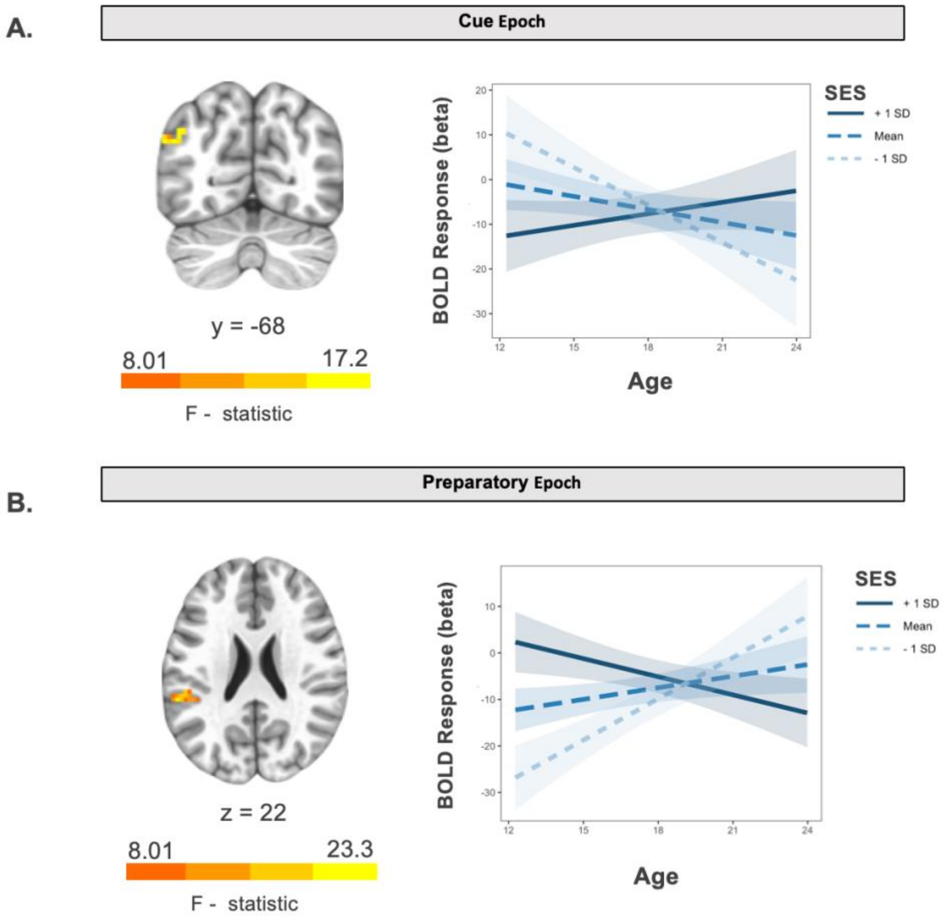


**Note.** Socioeconomic status (SES) by age interaction in the left angular gyrus during the cue epoch (Panel A). SES by age interaction in the left inferior parietal lobule during the preparatory epoch (Panel B). Voxelwise threshold $p < .005$, number of contiguous voxels > 20, $p < 0.05$ corrected.
